# Comparing approaches to quantify urbanization on a multicontinental scale

**DOI:** 10.1101/2024.04.30.591983

**Authors:** David Murray-Stoker, James S. Santangelo, Marta Szulkin, Marc T. J. Johnson

## Abstract

Urbanization is an increasingly prevalent driver of environmental, ecological, and evolutionary change in both terrestrial and aquatic systems, and it is important that our sampling designs accurately capture this urban environmental change. Common approaches to sampling urban environments include: urban-nonurban transects, which sample along urbanization gradients; random points, which sample locations at random within an area of interest; and systematic points, which sample locations based on a regularly-spaced grid from a predetermined starting point within the area of interest. Presently, we lack a comparative analysis of the efficacy of these different sampling designs in capturing variation in urban environments. Here, we compare the environmental variation captured by transect- and point-based sampling designs in 136 cities across six continents. We quantified and compared a common set of environmental variables for each sampling design, with variables capturing landcover, climate, and socioeconomic facets of urban environments. Mean landcover and socioeconomic metrics consistently differed among sampling designs, in contrast to climate variables that primarily varied among cities.

Additionally, changes in environmental variables with distance from the city centre depended on the sampling design, with this distance-by-sampling design interaction present in 27%–51% of cities, depending on the environmental variable. This implies that the rate of environmental change along urban-nonurban gradients frequently depends on the sampling design used. We also examined potential causes of deviations between transect- and point-based sampling designs and identified human population density and city area as common predictors of deviations between transect- and point-sampling designs. Our results show that sampling design can dictate how the urban environment is characterized, with sampling design as important − or more important − as the selected environmental variable. We further developed R code so researchers can implement these methods as they develop and validate sampling designs in novel or unstudied urban environments.

## Introduction

Anthropogenic disturbance is reshaping ecosystems across the globe (Vitousek et al. 1997, Malhi 2017), with urbanization being an increasingly prevalent driver of environmental change in both terrestrial and aquatic systems (Alberti et al. 2003, Grimm et al. 2008, Pickett et al. 2011). The majority of the human population resides within urban areas (United Nations Department of Economic and Social Affairs 2019), and the concomitant process of urbanization alters the landscape (Alberti et al. 2003, Grimm et al. 2008, Pickett et al. 2011). Urbanization is consistently associated with increased impervious surface cover, elevated pollution (e.g., light, noise, chemical), increased temperatures (i.e., urban heat island), and higher habitat fragmentation and degradation (Alberti et al. 2003, Grimm et al. 2008, Pickett et al. 2011).

Urban environmental changes alter community structure and ecosystem function (Grimm et al. 2008, Pickett et al. 2011, McFadden et al. 2023), including reduced biodiversity (McKinney 2008, Faeth et al. 2011, Piano et al. 2020), homogenized community composition (McKinney 2008, Faeth et al. 2011, Delgado-Baquerizo et al. 2021), and disrupted nutrient cycling (Grimm et al. 2008, Pickett et al. 2011, Ryan et al. 2022). There is also increasing evidence of urban-driven environmental change affecting evolution (Johnson and Munshi-South 2017, Szulkin et al. 2020, Diamond and Martin 2021), such as changes in natural selection (Caizergues et al. 2018, Irwin et al. 2018, Corsini et al. 2021, Santangelo et al. 2022), altered gene flow (Miles et al. 2018, Carlen and MunshiLJSouth 2021, Fusco et al. 2021), and increased mutation rates (Somers et al. 2002, Yauk et al. 2008). Notwithstanding the recent and rapid growth of empirical work in urban ecology and evolution, studies vary in their sampling designs, making it difficult to properly define and generalize patterns and processes.

Urban environments are studied using a variety of sampling designs. A common and effective approach in urban ecology and evolution is the use of urban-nonurban transects (McDonnell and Pickett 1990, McDonnell and Hahs 2008, Szulkin et al. 2020). In this design, spatial variation in urbanization is quantified by sampling from inside the city core and extending along a transect to suburban and rural areas (McDonnell and Pickett 1990, McDonnell and Hahs 2008, Szulkin et al. 2020); however, such urbanization gradients are not necessarily linear, as the effects of urbanization on the environment can be heterogeneous (McDonnell and Pickett 1990, Grimm et al. 2000, McDonnell and Hahs 2008, Szulkin et al. 2020, Alberti et al. 2020). Pairing urban and rural sites is a commonly-used alternative to the urban-nonurban transect (Winchell et al. 2016, Theodorou et al. 2016, Miles et al. 2022), but this design only captures categorical differences between habitats and cannot capture the spatial variation between sampling sites provided by the gradient approach. Random sampling within an area of interest can also be an effective design and is often preferred to accurately capture heterogeneity across a landscape (Bourdeau 1953, Kenkel et al. 1989, Albert et al. 2010). In contrast, systematic sampling, whereby sampling locations are selected based on a regularly-spaced grid from a predetermined starting point, can also be a useful sampling design to capture heterogeneous environments (Yates 1948, Iachan 1982, Kenkel et al. 1989). Logistical constraints related to environmental sampling can be non-trivial and affect which sampling approach is used. While urban-nonurban transects are the predominant method used in urban ecology and evolution (McDonnell and Hahs 2008, 2009, Szulkin et al. 2020), we lack a robust quantification and comparison of the efficacy of different sampling designs in capturing variation in the abiotic and biotic environment to guide best methodological practices.

In this study, we compared how different commonly implemented sampling designs captured urban environmental variation across a multicontinental sample of 136 cities sampled as part of the Global Urban Evolution (GLUE) project (Santangelo et al. 2022). We asked two research questions: (question 1) How are urban environments characterized by different sampling designs (i.e., urban-nonurban transects versus point sampling)? And (question 2) what underlies deviations in characterizations of the urban environment between sampling designs? To answer these questions, we developed tools to generate sampling designs within each city and then quantified a standardized suite of environmental variables (Table S1) for comparisons (question 1). We also calculated the deviation between sampling designs and used boosted regression trees to identify characteristics of cities that best predict deviations (question 2), with predictors related to city characteristics and urbanization intensity. Our objective was to understand how sampling designs differed in capturing urban environments, and to create a methodological resource in the R programming language for other urban ecology and evolution research projects.

## Methods

### Overview

We used 136 cities from the 160 cities in the GLUE project, which originally tested for evolutionary change in a plant defense along urbanization gradients (Santangelo et al. 2022). All selected cities had a minimum sampled transect length of ∼7.5 km. We used sites along the sampled transect for each city (hereafter GLUE transect) and we also generated sites using three additional sampling designs: (1) random transect direction, (2) random points, and (3) systematic points (Figure 1). We then extracted environmental variables (Table S1) for each site for all of the sampling designs (i.e., GLUE transect, random transect, random points, and systematic points; Figure 1).

**Figure 1:**
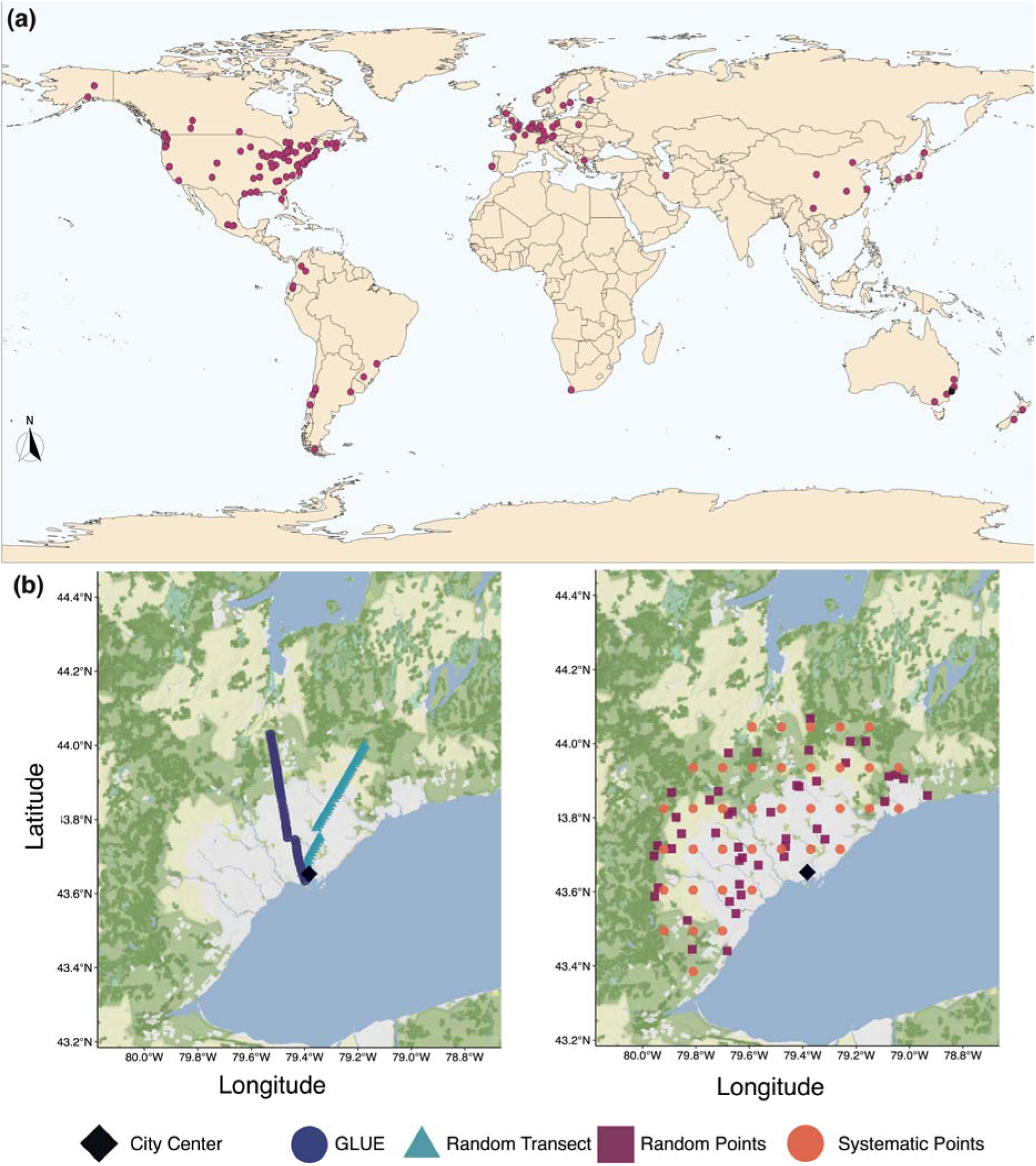
(a) Map of all 136 cities included in the analyses, with each city indicated with a pink circle. (b) Maps to visualize the different sampling designs. Using Toronto, Canada as an example, we display sites on the GLUE and random transects (left) and the random and systematic points (right). Sites are colored by sampling design: GLUE = dark blue circles, random transect = light blue triangles, random points = red squares, and systematic points = orange circles. The city centre is indicated by a black diamond. Sample sizes for each sampling design: GLUE = 43, random transect = 43, random points = 45, and systematic points = 37.

### Sampling Designs

#### GLUE transect

We used the urban-nonurban transect from each city for the GLUE project (Santangelo et al. 2022). Each of the GLUE transects was designed to (1) approximately comprise half urban and half rural habitats, (2) include a gradient of urban environmental change from the urban centre to outlying rural and/or natural habitats, and (3) avoid non-target environmental gradients (e.g., elevational gradients). As the length of each transect varied, the average distance between sampling sites was scaled to the total transect length, with a minimum distance of 200 m between sites (Santangelo et al. 2022). For each GLUE transect, we used the site GPS coordinates (latitude and longitude; coordinate reference system = EPSG 4326/WGS 84), and all GLUE transects retained their initial position and orientation (Santangelo et al. 2022).

#### Random transect

We generated a single, random transect for each city by rotating the GLUE transect by a random angle while keeping the urban centre of the transect fixed. Points classified over water were filtered from the dataset using a custom function relying on the ‘MODISTools‘ package (Tuck et al. 2014). To reduce sites positioned over large bodies of water (i.e., lakes, oceans), we manually set a range of random angles by which the GLUE transect could be rotated. Transects were rotated using the ‘rotate_2d()‘ function in the ‘rearrr‘ package (Olsen 2023).

#### Random points

We generated random points within the sampling area for each respective city. We set the sampling radius to the length from the city centre of the GLUE transect for a given city, and then randomly-sampled points within the resulting circle following a simple sequential inhibition (SSI) process using the ‘st_sample()‘ function with ‘type = “SSI”‘ in the ‘sf‘ (Pebesma et al. 2023a) and ‘spatstat‘ (Baddeley et al. 2023) packages. Using the SSI process allowed us to maintain a minimum, pairwise distance of 200 m between all points. We set the targeted number of random points equal to the number of sites sampled along the GLUE transect for each respective city. Targeted sample sizes were adjusted depending on the proportion of sites that were over water and therefore filtered from the dataset; sites were filtered by water cover type as described above.

#### Systematic points

We generated systematic points that followed grid-like sampling by making a rectangular sampling grid the length of the GLUE transect for each city using the ‘makegrid()‘ function in the ‘sp‘ package (Pebesma et al. 2023b). We again set the targeted number of systematic points equal to the number of sites sampled along the GLUE transect for the respective city, with sample sizes adjusted following the procedure for the random points. We then retained all sites within the circular sampling radius using a custom function relying on the ‘geospherè package (Hijmans 2019).

### Environmental Variables

We developed a method to quantify 19 environmental variables related to urbanization that spanned landcover, climate, and socioeconomic categories (Table S1). To illustrate the utility of our method, we focus on 8 variables described below. Distance from the city centre was calculated for each site as the distance on an ellipsoid using the ‘geospherè package (Hijmans 2019); we used the same coordinates for the city centres as the GLUE project (Santangelo et al. 2022). We estimated the impervious surface cover (ISC) using the 30-m resolution Global Man-Made Impervious Surface raster dataset (Brown de Colstoun et al. 2017), with ISC averaged within a 250-m buffer surrounding each site. We calculated the Human Influence Index (HII; estimate of human impacts) for each site using the 1-km resolution HII raster dataset (Wildlife Conservation Society - WCS and Center for International Earth Science Information Network - CIESIN - Columbia University 2005). We downloaded the Normalized Difference Vegetation Index (NDVI; quantifies the ‘greenness’ of vegetation) for each site across a 5-year period (1 January 2014-31 December 2018), which resulted in 115 measurements for each site (23 measurements per year). For each of the 5 years, we calculated the annual mean NDVI and then averaged these estimates to generate a single mean NDVI for each site; all NDVI measurements were calculated using the ‘MODISTools‘ package (Tuck et al. 2014). We calculated mean annual temperature and annual precipitation using the WorldClim database (Fick and Hijmans 2017) and 30 s resolution rasters. Finally, we used globally-gridded estimates for Shared Socioeconomic Pathway (SSP) projections for years 2030 and 2100 using 1-km resolution raster files (Wang and Sun 2022). Each SSP is a projection of gross domestic product based on a series of demographic, economic, and political development variables under global climate change, and each SSP has different constraints and assumptions for the projections (Dellink et al. 2017, O’Neill et al. 2017). We used SSP2 (middle of the road) as a heuristic, whereby this projection assumes current social and economic trends continue and do not strongly deviate from historical patterns, although with lower increases in urbanization coupled with lower improvements to human development (Dellink et al. 2017, O’Neill et al. 2017). Values for environmental variables calculated from raster files (i.e., ISC, HII, climate variables, and socioeconomic variables) relied on custom functions using the ‘raster‘ package (Hijmans et al. 2023). Focal environmental variables were categorized as: landcover = distance, ISC, HII, and mean NDVI; climate = mean annual temperature and annual precipitation; and socioeconomic = SSP2 2030 and SSP2 2100. A full description of how all landcover, climate, and socioeconomic variables were calculated is available in the Supplementary Methods (Appendix S1).

### Statistical Analyses

#### Comparison of sampling sites

To determine if the number of sampling sites differed among sampling designs aftering adjusting the target sample sizes and filtering points over water, we used a one-way analysis of variance (ANOVA). For each city, we calculated the number of sites for each sampling design. We then compared the number of sites by sampling design by fitting the model as:

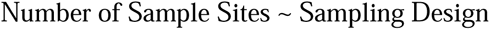

Model assumptions were evaluated using the ‘performancè package (Lüdecke et al. 2021), and the F-test was conducted with Type I sums-of-squares. We calculated effect size for sampling design as eta-squared (η^2^), which represents the proportion of variance explained by that model term (Cohen 1988). We found that the number of sampling sites was similar across all sampling designs (F_3, 540_ = 0.446, P = 0.720, η^2^ = 0.002).

#### Environmental variable-by-distance linear mixed-effects models

To determine if sampling designs differentially-captured environmental change along urbanization gradients, we used linear mixed-effects models (LMMs). We first calculated a standardized distance, whereby distance from the city centre was standardized between 0 and 1, where 0 = city centre and 1 = most distant non-urban sampling site along the transect. By standardizing distance, all points and transects were on the same scale, which allowed for direct comparisons. All LMMs were fitted as:

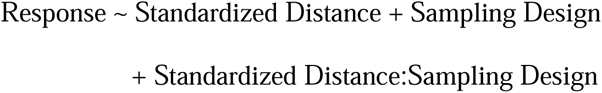

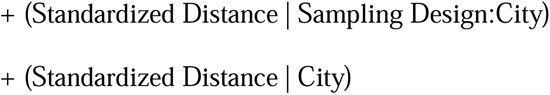

where the response was one of the remaining 7 focal environmental variables (i.e., distance was removed as a response and only used as a predictor) and standardized distance, sampling design, and the interaction were fixed effects. Responses along the urbanization gradients were allowed to vary by city (i.e., random slope and intercept for each city) and the interaction between sampling design and city (i.e., random slope for each level of the sampling design-by-city interaction). All LMMs were fitted with the bobyqa optimizer using the ‘lme4‘ (Bates et al. 2015) and ‘lmerTest‘ (Kuznetsova et al. 2017) packages. Estimated marginal means of linear trends (i.e., slopes) were calculated using ‘emtrends()‘ in the ‘emmeans‘ package (Lenth et al. 2022).

We calculated F-tests of fixed effects using Type III sums-of-squares, with denominator degrees-of-freedom estimated using the Satterthwaite method (Satterthwaite 1946). Model assumptions were evaluated using the ‘performancè package (Lüdecke et al. 2021). We calculated effect sizes for fixed effects as partial eta-squared (η^2^_P_), while effect sizes for the random effects were quantified as the intraclass correlation coefficient (ICC), or the proportion of variance explained by the random grouping factor (Nakagawa et al. 2017). We also quantified the variation explained by fixed effects (marginal R^2^, R^2^_Marginal_) and the combined fixed and random effects (conditional R^2^, R^2^, R^2^_Conditional_) using the ‘r2_nakagawà function in the ‘performancè package (Nakagawa et al. 2017, Lüdecke et al. 2021).

#### Environmental Heterogeneity ANOVA

To compare the environmental heterogeneity captured by the different sampling designs, we used ANOVA. To quantify within-city environment heterogeneity, we first performed a principal components analysis (PCA) of all 8 focal environmental variables (Table S1) for each city-by-sampling-design combination. This was done by standardizing the environmental variables (i.e., mean = 0, standard deviation = 1) and transforming the variables into a Euclidean distance matrix. We quantified environmental heterogeneity as the Euclidean distance from each sampling location to the centroid, with separate centroids for each city and sampling design. Environmental heterogeneity was quantified using the ‘betadisper()‘ function in the ‘vegan‘ package (Oksanen et al. 2020). We fitted the linear mixed-effects model as:

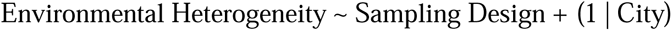

where the response was environmental heterogeneity, sampling design was the fixed effect, and city was fitted as a random intercept. The LMM was fitted using the ‘lme4‘ (Bates et al. 2015) and ‘lmerTest‘ (Kuznetsova et al. 2017) packages. We calculated the F-test using Type III sums-of-squares, with denominator degrees-of-freedom estimated using the Satterthwaite method (Satterthwaite 1946). Model assumptions were evaluated using the ‘performancè package (Lüdecke et al. 2021). We calculated the effect size for sampling design as eta-squared (η^2^), while the effect size for the random intercept of city was quantified as the intraclass correlation coefficient (ICC). Post-hoc comparisons were conducted using estimated marginal means weighted by cell frequencies using the ‘emmeans‘ package (Lenth et al. 2022).

#### City-specific regressions

To further evaluate changes in environments along urbanization gradients and specifically the frequency of the interaction between sampling design and distance, we fit individual linear regressions for each of the cities. We followed a similar procedure for the city-specific regressions as we did for the environmental variable-by-distance LMMs. For each city, linear models were fitted with the form:

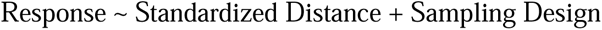

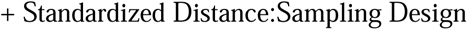

where the response was one of the 7 environmental variables and standardized distance, sampling design, and the interaction were the predictors. We calculated the ANOVA for each linear model using Type III sums-of-squares using the ‘Anova()‘ function in the ‘car‘ package (Fox and Weisberg 2018), with effect sizes calculated as partial eta-squared (η^2^_P_).

#### Deviation predictor boosted regression trees

We used stochastic boosted regression trees (BRTs) to identify predictors of deviations in the ability of transect- and point-sampling designs to capture environmental change along urbanization gradients in each city (Elith et al. 2008, Pichler and Hartig 2023). We used BRTs because they are not restricted by assumptions of parametric methods, such as assumptions of normality, independence between predictor variables, or linear relationships between predictor and response variables (Elith et al. 2008, Pichler and Hartig 2023).

For each environmental variable, we calculated the mean slope for the transects (GLUE and random transects) and the mean slope for the points (random and systematic points). Transects and points were combined into respective categories because the different sampling designs within each of these categories yielded quantitatively and qualitatively similar results across analyses and environmental variables, which allowed us to reduce the number of comparisons to be made in the analyses. We calculated the deviation between sampling designs as the absolute difference between the transects slope and the points slope (i.e., Deviation = Transects Slope – Points Slope) for all 136 cities. Slopes were extracted from the environmental variable-by-distance LMMs using ‘ggpredict()‘ in the ‘ggeffects‘ package (Lüdecke 2018).

We fitted separate BRTs for each of the environmental variables, with the deviation associated with the environmental variable as the response and city area, human population size, human population density, city age, and the number of nearby cities as predictor variables (variables compiled by Santangelo et al. 2022). Model parameters for each BRT were selected based on model training using the ‘train()‘ function in the ‘caret‘ package (Kuhn et al. 2023). Individual BRTs varied in the optimal number of trees, interaction depth, learning rate, and minimum observations per node. All BRTs used ten-fold cross validation and stochastic boosting with a bagging fraction of 50% (Elith et al. 2008). Effects of a predictor on the deviation between transect and point slopes were quantified as relative influence, with the relative influence within a BRT scaled between 0 (low influence) and 100 (high influence). We fitted all BRTs using the ‘gbm()‘ function in the ‘gbm‘ package (Greenwell et al. 2020).

All above analyses were performed using R (version 4.2.3; R Core Team 2023) in the RStudio environment (version 2023.09.1; RStudio Team 2022). Data management and figure creation were facilitated using the ‘tidyversè (Wickham et al. 2019) and ‘ggpubr‘ (Kassambara 2020) packages. Effect sizes were quantified using the ‘effectsizè (η^2^and η^2^_P_; Ben-Shachar et al. 2020) and ‘performancè (ICC; Lüdecke et al. 2021) packages.

## Results

### Variation in the Urban Environment by Distance and Sampling Design

Environmental variables varied by distance, sampling design, and the interaction (Table 1, Figures 2–3 and S1–S4). Both distance and sampling design had consistent effects across all environmental variables, although effects were stronger for landcover variables (distance η^2^_P_= 0.235−0.903, sampling design η^2^_P_= 0.015−0.408; Table 1, Figures 2 and 3) and socioeconomic variables (distance η^2^ = 0.915−0.919, sampling design η^2^_P_= 0.222−0.237; Table 1, Figures S3 and S4) than for climate variables (distance η^2^_P_= 0.009−0.125; sampling design η^2^_P_ = 0.004−0.007; Table 1, Figures 2 and 3). For example, GLUE transects had 16% higher city-wide mean estimates ISC and 5% higher HII compared to random transects. In contrast, GLUE transects had 32−33% higher city-wide mean estimates of ISC and 5% higher estimates of HII than both random and systematic point sampling designs; random transects followed similar patterns (Table 1, Figure 2). Mean values of climate variables never differed by more than ∼1.6% among sampling designs (Table 1, Figure 2). City-wide mean estimates of socioeconomic variables (i.e., SSP2 2030 and SSP2 2100) were higher for GLUE transects compared to random transects (10% higher), random points (12% higher), and systematic points (12% higher; Figure S3), while random transects were similar to point-based designs (percent difference = 2−3%; Figure S3); random and systematic points had similar estimates of socioeconomic variables (< 1% difference; Figure S3).

**Figure 2:**
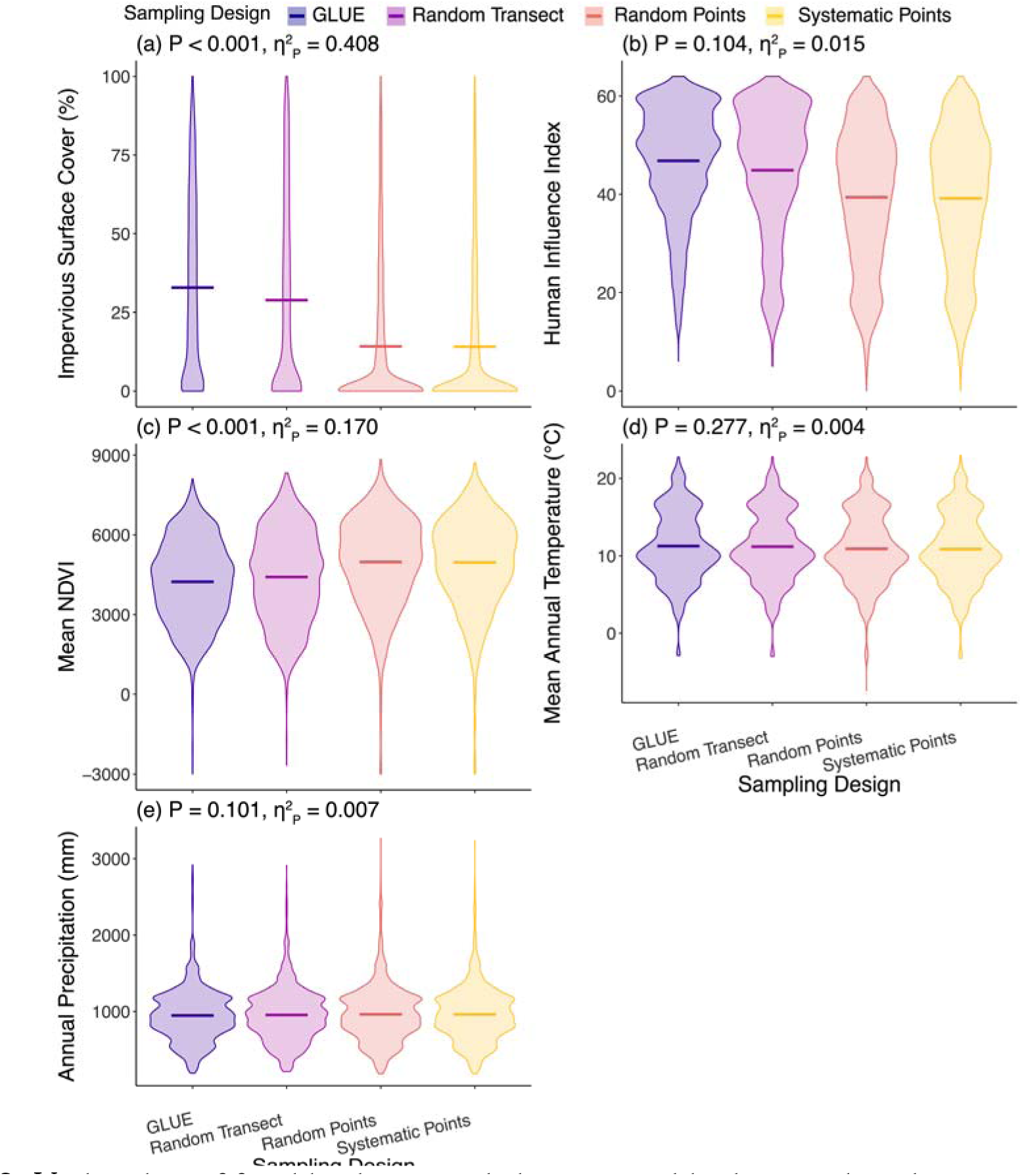
Violin plots of focal landcover and climate variables by sampling design across all cities. Violins are colored by sampling design, where GLUE transect = dark blue, random transect = purple, random points = orange, and systematic points = yellow. Crossbars indicate the mean across cities. Inset text provides the P-value and effect size (partial eta-squared, η^2^_P_) for the fixed effect of sampling design. Detailed statistics are provided in Table 1.

**Figure 3:**
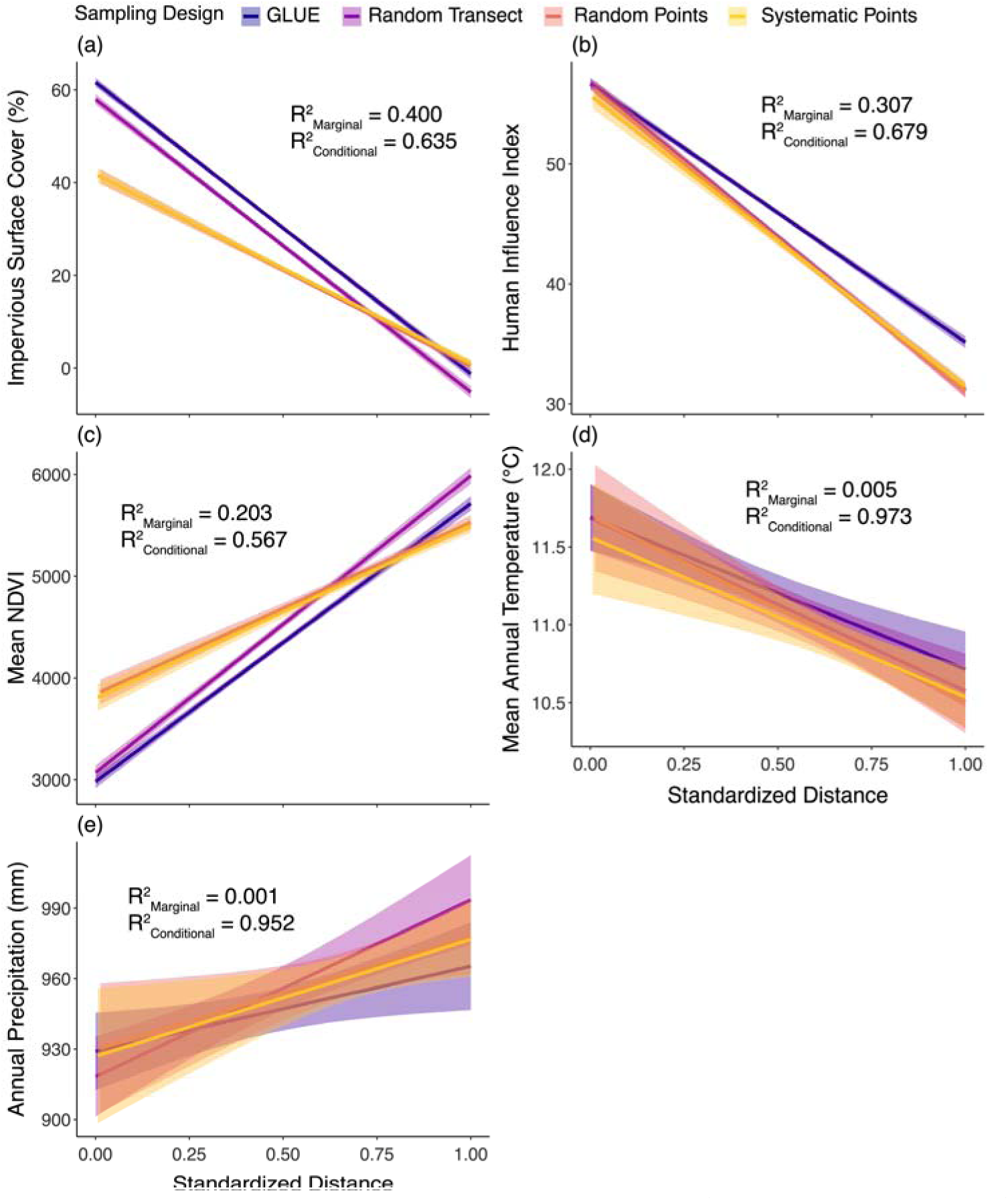
Relationships between focal environmental variables (landcover and climate) and standardized distance to the city center (0 = city center, 1 = rural limit) by sampling design. Lines are lines-of-best fit (± 95% confidence interval) and are colored by sampling design: GLUE transect = dark blue, random transect = purple, random points = orange, and systematic points = yellow. We report the marginal R^2^ (variance explained by fixed effects, R^2^_Marginal_) and conditional R^2^ (variance explained by fixed and random effects, R^2^_Conditional_). Detailed test statistics provided in Table 1.

**Table 1:** Summary of the environmental variable-by-distance linear mixed-effects models. For each model term [distance, sampling design, and the interaction (distance X sampling design)], we report the P-value and effect size (partial eta-squared, η^2^_P_). We also report the marginal and conditional R^2^ values for each model. Variables are abbreviated as follows: ISC = impervious surface cover, HII = Human Influence Index, NDVI = normalized difference vegetation index, GDP = gross domestic product, and SSP = shared socioeconomic pathway.

| Environmental Variable | Distance | | Sampling Design | | Distance × Sampling Design | | $R^2$ | |
| --- | --- | --- | --- | --- | --- | --- | --- | --- |
| | P-value | $\eta^2_p$ | P-value | $\eta^2_p$ | P-value | $\eta^2_p$ | Marginal | Conditional |
| **Landcover** |  |  |  |  |  |  |  |  |
| **ISC** | < 0.001 | 0.903 | < 0.001 | 0.408 | < 0.001 | 0.352 | 0.400 | 0.635 |
| **HII** | < 0.001 | 0.872 | 0.104 | 0.015 | < 0.001 | 0.044 | 0.307 | 0.679 |
| **Mean NDVI** | < 0.001 | 0.235 | < 0.001 | 0.170 | < 0.001 | 0.020 | 0.203 | 0.567 |
| Min NDVI | < 0.001 | 0.073 | < 0.001 | 0.041 | < 0.001 | 0.008 | 0.038 | 0.671 |
| Max NDVI | < 0.001 | 0.318 | < 0.001 | 0.228 | < 0.001 | 0.024 | 0.300 | 0.589 |
| **Climate** |  |  |  |  |  |  |  |  |
| **Mean Annual Temperature** | < 0.001 | 0.125 | 0.277 | 0.004 | < 0.001 | 0.001 | 0.005 | 0.973 |
| Temperature Seasonality | 0.001 | 0.077 | < 0.001 | 0.066 | 0.448 | 0.006 | < 0.001 | 0.998 |
| Range Annual Temperature | 0.759 | 0.001 | 0.125 | 0.015 | 0.235 | 0.010 | < 0.001 | 0.995 |
| **Annual Precipitation** | < 0.001 | 0.009 | 0.101 | 0.007 | < 0.001 | 0.005 | 0.001 | 0.952 |
| Precipitation Seasonality | < 0.001 | 0.002 | 0.983 | < 0.001 | 0.547 | < 0.001 | < 0.001 | 0.992 |
| Aridity Index | < 0.001 | 0.042 | 0.225 | 0.003 | < 0.001 | 0.002 | 0.007 | 0.894 |
| **Socioeconomic** |  |  |  |  |  |  |  |  |
| GDP 2005 | < 0.001 | 0.905 | < 0.001 | 0.168 | < 0.001 | 0.042 | 0.356 | 0.650 |
| SSP1 2030 | < 0.001 | 0.919 | < 0.001 | 0.236 | < 0.001 | 0.076 | 0.355 | 0.600 |
| SSP1 2100 | < 0.001 | 0.914 | < 0.001 | 0.220 | < 0.001 | 0.073 | 0.349 | 0.593 |
| **SSP2 2030** | < 0.001 | 0.919 | < 0.001 | 0.237 | < 0.001 | 0.077 | 0.355 | 0.600 |
| **SSP2 2100** | < 0.001 | 0.915 | < 0.001 | 0.222 | < 0.001 | 0.073 | 0.350 | 0.594 |
| SSP5 2030 | < 0.001 | 0.353 | < 0.001 | 0.083 | < 0.001 | 0.005 | 0.361 | 0.562 |
| SSP5 2100 | < 0.001 | 0.910 | < 0.001 | 0.210 | < 0.001 | 0.073 | 0.338 | 0.582 |

Note: Focal environmental variables are in bold.

We also frequently identified an interaction between distance and sampling design, which indicates that changes in environmental variables with distance depended on the sampling design (Table 1, Figures 3 and S3). For example, both GLUE and random transects had higher estimates of ISC than random or systematic points at lower distances, but random and systematic points had higher estimates of ISC as distance from the city centre increased (Table 1, Figures 3). In other words, point sampling captured less impervious surface cover in the city centre than transect sampling (Table 1, Figures 3). Patterns were quantitatively and qualitatively consistent for the non-focal landcover and climate variables (Table 1, Figures S1 and S2) and the non-focal socioeconomic variables (Table 1, Figures S3 and S4).

When decomposed using city-specific regressions, the interaction between distance and sampling design was common (prevalence = 27%−51% of cities; Table 2, Figure 4). For example, there was a negative relationship between ISC and standardized distance from the city centre across all sampling designs, but the GLUE and random transect slopes were more negative than the random or systematic point slopes [GLUE (β ± standard error) = −63.99 ± 1.97, random transect = −64.40 ± 1.97, random points = −41.07 ± 2.05, systematic points = −40.38 ± 2.05; Figure 3]. In contrast, the relationship between HII and standardized distance from the city centre was similar across all sampling designs (GLUE = −21.83 ± 1.05, random transect = −26.22 ± 1.05, random points = −25.82 ± 1.08, systematic points = −24.08 ± 1.08; Figure 3). Similarly, relationships between mean annual temperature and distance from the city centre were negative across all sampling designs (GLUE = −0.917 ± 0.034, random transect = −1.126 ± 0.034, random points = −1.122 ± 0.42, systematic points = −1.058 ± 0.043; Figure 3).

**Figure 4:**
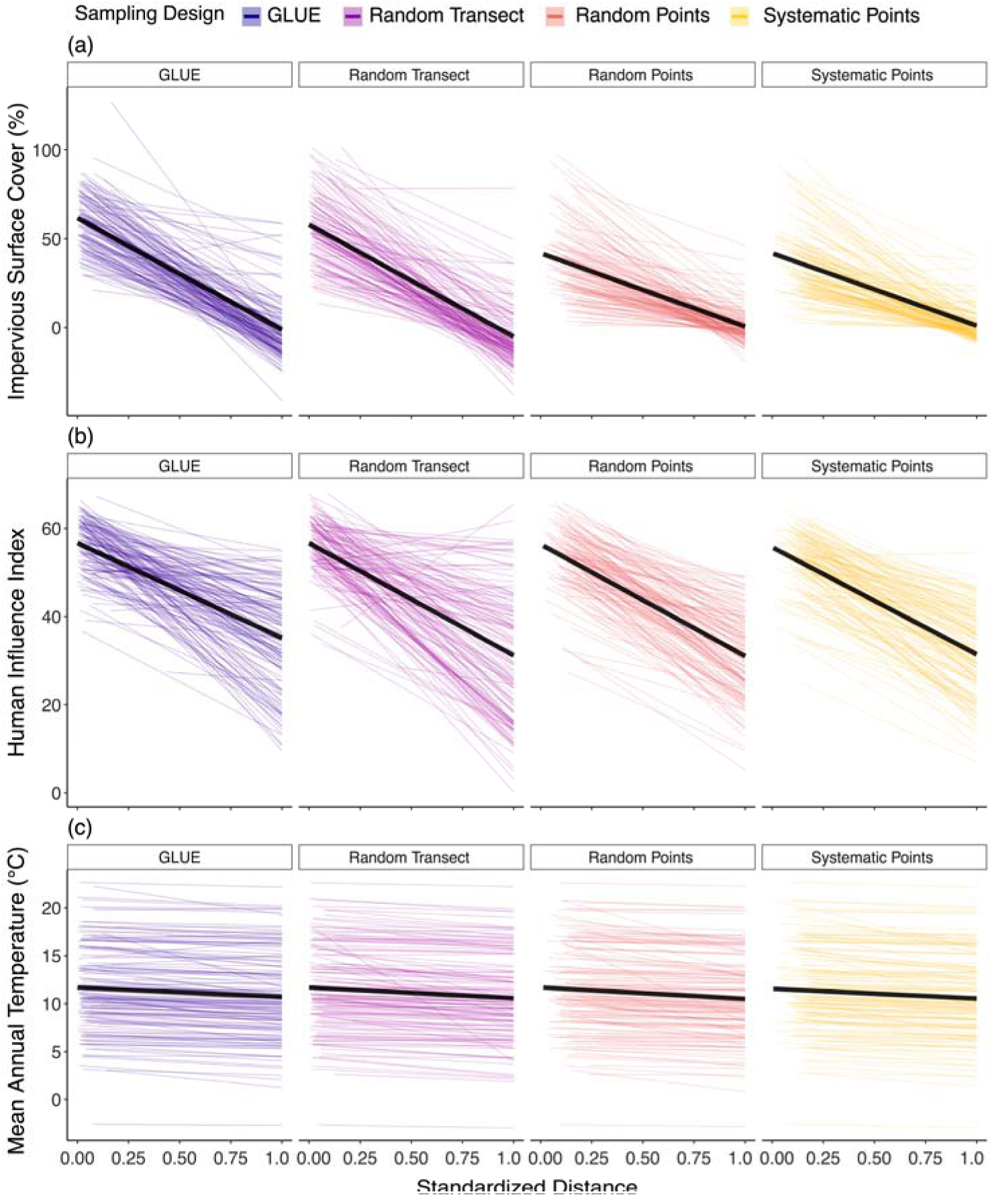
Relationships between environmental variables (impervious surface cover, Human Influence Index, and mean annual temperature) and standardized distance to the city center (0 = city center, 1 = nonurban limit) by sampling design. Lines are lines-of-best fit (± 95% confidence interval) and are colored by sampling design: GLUE transect = dark blue, random transect = purple, random points = orange, and systematic points = yellow. Each colored line represents the relationship for an individual city, with the larger black line indicating the mean slope across all cities. Summary statistics are provided in Table 2.

**Table 2:** Summary of the city-specific regressions for each environmental variable For each model term [distance, sampling design, and the interaction (distance X sampling design)], we report the percent of models (N = 136 cities) where the term had a P-value < 0.001 (% < 0.001), the mean P-value (standard error), and the mean η^2^_P_ (standard error). Variables are abbreviated as follows: ISC = impervious surface cover, HII = Human Influence Index, NDVI = normalized difference vegetation index, GDP = gross domestic product, and SSP = shared socioeconomic pathway.

| Environmental Variable | Distance | | | Sampling Design | | | Distance × Sampling Design | | |
| --- | --- | --- | --- | --- | --- | --- | --- | --- | --- |
| | % $< 0.001$ | P (SE) | $\eta^2_P$ (SE) | % $< 0.001$ | P (SE) | $\eta^2_P$ (SE) | % $< 0.001$ | P (SE) | $\eta^2_P$ (SE) |
| **ISC** | 90.441 | 0.021 (0.009) | 0.523 (0.012) | 52.206 | 0.073 (0.015) | 0.114 (0.009) | 38.971 | 0.122 (0.020) | 0.102 (0.007) |
| **HII** | 69.118 | 0.065 (0.015) | 0.489 (0.015) | 5.147 | 0.347 (0.027) | 0.141 (0.010) | 29.412 | 0.145 (0.020) | 0.089 (0.006) |
| **Mean NDVI** | 79.265 | 0.041 (0.011) | 0.352 (0.015) | 33.088 | 0.125 (0.018) | 0.092 (0.008) | 27.206 | 0.145 (0.020) | 0.077 (0.005) |
| **Mean Annual Temperature** | 74.265 | 0.066 (0.016) | 0.505 (0.019) | 5.882 | 0.323 (0.027) | 0.201 (0.013) | 41.176 | 0.131 (0.020) | 0.107 (0.008) |
| Temperature Seasonality | 55.882 | 0.115 (0.020) | 0.291 (0.023) | 6.618 | 0.344 (0.027) | 0.169 (0.012) | 32.353 | 0.171 (0.024) | 0.089 (0.006) |
| Range Annual Temperature | 54.412 | 0.116 (0.021) | 0.265 (0.022) | 10.294 | 0.348 (0.028) | 0.173 (0.11) | 30.147 | 0.157 (0.022) | 0.081 (0.005) |
| **Annual Precipitation** | 45.588 | 0.114 (0.020) | 0.161 (0.014) | 5.882 | 0.416 (0.029) | 0.197 (0.012) | 41.912 | 0.121 (0.020) | 0.105 (0.008) |
| Precipitation Seasonality | 48.529 | 0.123 (0.020) | 0.181 (0.017) | 3.676 | 0.451 (0.028) | 0.174 (0.011) | 30.882 | 0.132 (0.020) | 0.092 (0.006) |
| Aridity Index | 47.059 | 0.129 (0.021) | 0.280 (0.021) | 9.559 | 0.370 (0.029) | 0.180 (0.013) | 35.294 | 0.147 (0.022) | 0.098 (0.007) |
| GDP 2005 | 94.118 | 0.009 (0.007) | 0.431 (0.013) | 56.618 | 0.045 (0.011) | 0.065 (0.006) | 43.382 | 0.061 (0.013) | 0.107 (0.006) |
| SSP1 2030 | 90.441 | 0.012 (0.007) | 0.357 (0.012) | 60.294 | 0.042 (0.012) | 0.057 (0.006) | 50.735 | 0.047 (0.010) | 0.114 (0.006) |
| SSP1 2100 | 90.441 | 0.012 (0.007) | 0.357 (0.012) | 60.294 | 0.041 (0.012) | 0.057 (0.006) | 51.471 | 0.046 (0.010) | 0.114 (0.006) |
| **SSP2 2030** | 90.441 | 0.012 (0.007) | 0.357 (0.012) | 60.294 | 0.042 (0.012) | 0.057 (0.006) | 50.735 | 0.047 (0.010) | 0.114 (0.006) |
| **SSP2 2100** | 90.441 | 0.012 (0.007) | 0.357 (0.012) | 60.294 | 0.041 (0.012) | 0.057 (0.006) | 50.735 | 0.046 (0.010) | 0.114 (0.006) |
| SSP5 2030 | 90.441 | 0.012 (0.007) | 0.357 (0.012) | 60.294 | 0.042 (0.012) | 0.057 (0.006) | 50.735 | 0.047 (0.010) | 0.114 (0.006) |
| SSP5 2100 | 90.441 | 0.013 (0.008) | 0.357 (0.012) | 60.294 | 0.041 (0.012) | 0.057 (0.006) | 51.471 | 0.046 (0.010) | 0.114 (0.006) |

Note: Focal variables are in bold. Additionally, minimum and maximum NDVI were not fitted as the variance component in the ANOVA was effectively 0.

### Environmental Heterogeneity

Environmental heterogeneity varied considerably by sampling design (F_3, 21065.71_ = 404.244, P < 0.001, η^2^ = 0.054, ICC = 0.223; Figure S5). Compared to random and systematic point sampling designs, GLUE and random transects captured 96-102% more heterogeneity (Figure S5). Random transects had higher environmental heterogeneity than GLUE transects (random transect = 7% higher; Figure S5), but there were no discernible differences between random and systematic points (systematic points = 1% higher; Figure S5).

### Effects of City Characteristics on Deviations Between Sampling Designs

Human population density and city area were the primary predictors of deviation in slope between transect- and point-sampling designs, but these results depended on the specific environmental variable (Figure 5). Impervious surface cover deviations were primarily influenced by human population density (relative influence (RI) = 28.72), with city area of similar influence (RI = 26.18). Similarly, HII deviations were primarily influenced by human population density (RI = 56.43), with city area also having a marked effect (RI = 20.37). Mean annual temperature was primarily influenced by human population density (RI = 78.09). Human population size was of primary influence for SSP2 2030 (RI = 54.89) and SSP2 2100 (RI = 68.94) with city area of secondary influence (SSP2 2030 RI = 31.50, SSP2 2100 RI = 14.20, Figure S6). Patterns of deviation predictors were qualitatively similar for non-focal landcover, climate, and socioeconomic variables (Figure S6).

**Figure 5:**
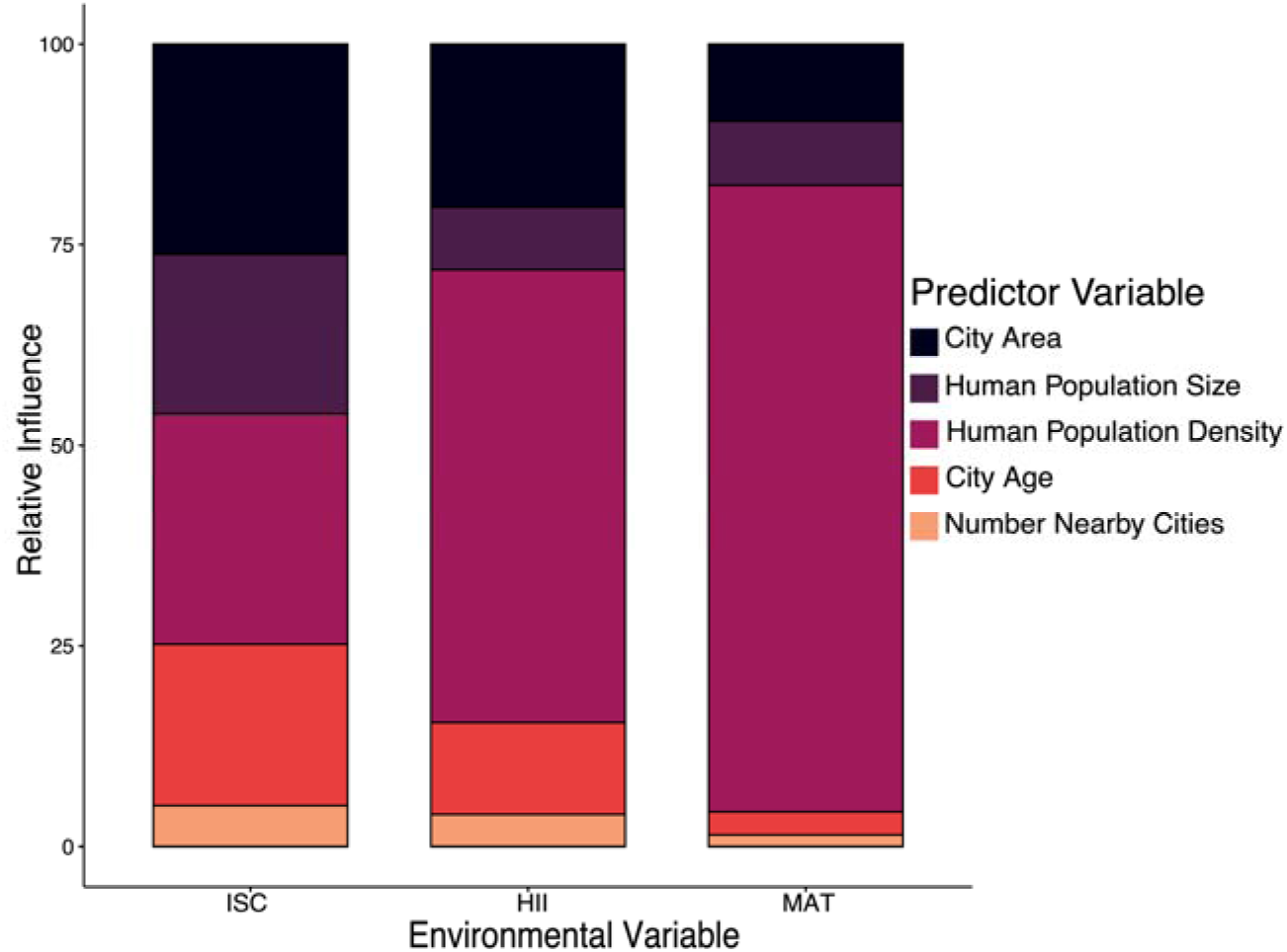
Relative influence of city characteristics (city area, human population size, human population density, city age, and number of nearby cities) for predicting deviations between transects (GLUE and random transects) and points (random and systematic points) for impervious surface cover (ISC), Human Influence Index (HII), and mean annual temperature (MAT).

## Discussion

Our comparative analyses of a large sample of cities across continents demonstrate that characterizations of urban environments depend on sampling design. We found that mean values of landcover and socioeconomic metrics consistently varied by sampling design, in contrast to climate variables that primarily varied by city (question 1). Specifically, sites along urban-nonurban transects captured greater estimates of environmental features typically linked with urbanization and associated economic activity (e.g., impervious surface cover; Figure 2). Moreover, change in the environment with distance from the city centre varied by sampling design (Figure 3), suggesting that sampling designs differentially sample environmental change along urban-nonurban gradients. Deviations between transect and point sampling designs depended on the environmental variable (question 2), with city area and human population density being common predictors of deviations between sampling designs (Figure 5). Below, we discuss the implications and applications of our work within the broader context of urban ecology.

### Sampling designs can differentially characterize urban environments

Studies in urban environments commonly use both transect- and point-based sampling designs, and our results show that the sampling design chosen can dictate how the urban environment is characterized. Importantly, while there were minor differences in landcover and socioeconomic variables between GLUE and random transects, our results indicate that sampling a different transect had less of an influence than the broader effect of using transect-versus point-sampling designs. Transects sampled more of the urban core, as indicated by increased estimates of distance from the city centre for point-based designs and increased impervious surface cover for transect-based designs (Figure 3). Additionally, point sampling designs had considerably lower estimates of GDP compared to GLUE and transects, and this trend was consistent for all shared socioeconomic pathways. As expected, climate variables differed almost exclusively by city identity and not sampling design, because cities were distributed at the multi-continental scale and situated within different regional climates (Jiang et al. 2017). We still observed consistent trends of increased temperatures in the urban core (i.e., within cities), which is consistent with the urban heat island effect (Oke 1973, Peng et al. 2012), and all sampling designs tended to capture changes in temperature in a similar way.

We found that city area was a frequent predictor of deviations between sampling designs. There was a wide range in city area (mean ± standard error = 874.6 km^2^ ± 117.6 km^2^, min = 8.81 km^2^, max = 9212.1 km^2^), but the number of sampling sites was similar across all sampling designs. As the purpose of our study was to compare sampling designs, we constrained the number of sites so sample sizes would be relatively equal across sampling designs. Given the wide range in city area and relatively constant sample sizes, it is likely that urban areas – particularly in larger cities – were undersampled compared to periurban and rural areas. Random points had the potential to be distributed across the entire sampling radius, and systematic points covered the sampling radius with an equally-spaced grid, with greater spacing between points as the sampling radius increased. In contrast, GLUE and random transects followed an approximately linear path from the urban core, with regularly-spaced intervals between sampling locations (Santangelo et al. 2022). Moreover, GLUE transects were designed so half of the transect represented urban and suburban areas and the other half represented periurban and rural areas (Santangelo et al. 2022), which could explain why environmental features differed between transect and point samples. Assuming the gradient is evenly divided between urban and nonurban, a larger portion of the total sampling area will necessarily be in periurban and rural areas than the urban centre. Point sampling designs could mitigate this issue by increasing the number of sites within urban areas.

Because cities are not necessarily monocentric but instead a mosaic of biological, chemical, and physical components (McDonnell and Pickett 1990, Pickett et al. 1997, 2011, Grimm et al. 2000, 2008, Szulkin et al. 2020), random and systematic points have the potential to differentially capture the heterogeneity in the environment because they have a wider distribution than transects. Surprisingly, we found that transect designs captured more environmental heterogeneity than either random or systematic points. These results suggest that, although sample sizes could have led to undersampling in larger cities, transect sampling designs are best suited for capturing environmental heterogeneity along urban-rural gradients.

### Applications to urban ecology and evolution

Urban ecology and evolution research is a rapidly-growing field (Alberti et al. 2003, Pickett et al. 2011, Johnson and Munshi-South 2017, Szulkin et al. 2020, Diamond and Martin 2021), and we have established a method for researchers to develop and validate sampling designs in urban environments. One application of this method is to plan the design before any sampling has occurred. Another application is to compare sampling designs, which allow researchers to determine if there are aspects of the urban environment that are over- or under-represented by the sampled locations. Finally, by including projections of shared socioeconomic pathways, researchers can use our method to establish sampling designs to evaluate the effects of socioeconomic and environmental change through time (Ramalho and Hobbs 2012, Dellink et al. 2017, O’Neill et al. 2017). In order to apply our R-based method to other urban ecology and evolution projects, researchers only need to have site coordinates for a transect- or point-based sampling design, download the necessary raster files (Wildlife Conservation Society - WCS and Center for International Earth Science Information Network - CIESIN - Columbia University 2005, Brown de Colstoun et al. 2017, Fick and Hijmans 2017, CGIAR-CSI 2019, Wang and Sun 2022, Zomer et al. 2022), and use the R code that has been provided on Zotero (Murray-Stoker et al. 2024). Extensive annotations allow for researchers to use and modify the code as needed for their projects.

### Limitations and considerations

We have provided an extensive comparison of sampling designs within urban environments; however, there are some limitations and considerations with our study. First, our study was focused on comparing sampling designs, not addressing how urbanization can be quantified. The methods used and variables selected for quantifying urbanization remain a subject of debate (McDonnell and Hahs 2008, 2013, Moll et al. 2019, Szulkin et al. 2020), but to make comparisons at the multi-continental scale we had to use broader variables. Given the differences we observed among sampling designs using the same environmental variables, our results suggest that sampling design could at least be as important − if not more important − as the choice of environmental variables, particularly for landcover and socioeconomic variables. Second, notwithstanding differences between transect- and point-based sampling designs, the choice of sampling design will be dictated by the research objectives and study organisms (Kenkel et al. 1989, Szulkin et al. 2020). When aiming to quantify urban environments, we encourage the use of transect methods when possible because they capture more environmental heterogeneity and are more easily and efficiently adjusted to equally capture urban and nonurban environments. We also acknowledge this will not always be possible or desirable for all projects (e.g., distribution of study organisms, logistical constraints). For example, paired urban-nonurban sampling may be a valuable alternative for genomic approaches to explain urban-nonurban phenotypic differences (e.g., Lotterhos and Whitlock 2015). Alternatively, dyads of contrasting environments typical of the urban mosaic (e.g., urban park and residential area) can be sampled at similar distances from the city centre to infer the impact of urbanization while controlling for spatial variation in urban environmental change (e.g., Corsini et al. 2019).

Finally, our method is currently restricted to use in terrestrial ecosystems. River and stream ecosystems often follow a dendritic network (Campbell Grant et al. 2007, Altermatt 2013), which requires longitudinal sampling to identify the source of anthropogenic disturbances and evaluate the localized impact of the (non)local disturbances on communities and ecosystems. In addition to rivers and streams, lentic freshwater (e.g., natural ponds, stormwater ponds, and wetlands) and coastal marine ecosystems can integrate the effects of urbanization from a large drainage area (Campbell Grant et al. 2007, Alter et al. 2021). Despite these limitations and considerations, we contend that our approach provides a robust comparison of sampling designs and allows for other researchers to apply the method to projects in urban ecology and evolution.

## Conclusions

Our study presents a comparative quantification of the efficacy of commonly used sampling designs in capturing urban-nonurban environmental change at a multi-continental scale. We found that sampling designs differentially-characterized urban environments, with urban-nonurban transects sampling a greater proportion of the urban area and having more environmental heterogeneity than random or systematic point designs. Landcover and socioeconomic variables varied along urbanization gradients and among sampling designs, while climate variables depended on the city. We also identified city area and human population as common predictors of deviations between sampling designs. As urban environments continue to expand and more people inhabit cities (Grimm et al. 2008, Pickett et al. 2011, United Nations Department of Economic and Social Affairs 2019), it is imperative to understand how these environmental changes affect ecology and evolution in urban ecosystems. We aim for our method to be applied to this challenge by designing robust studies that capture the ecological and evolutionary consequences of urbanization.

## Acknowledgements

We thank all the researchers who participated in the Global Urban Evolution Project, which facilitated the current project. M. Szulkin was funded with grants Sonata BIS 2014/14/E/NZ8/00386 and OPUS 2021/41/B/NZ8/04472. M. Johnson was funded by an NSERC Discovery Grant, Canada Research Chair, and E.W.R. Steacie Fellowship.

## Author Contributions

**David Murray-Stoker:** Conceptualization (equal); Methodology (equal); Data curation (lead); Formal analysis (lead); Software (lead); Investigation (lead); Visualization (lead); Writing – original draft (lead); Writing – review and editing (equal)

**James S. Santangelo:** Conceptualization (equal); Methodology (equal); Data curation (supporting); Formal analysis (supporting); Software (supporting); Writing – review and editing (equal)

**Marta Szulkin:** Conceptualization (equal); Methodology (equal); Data curation (supporting); Formal analysis (supporting); Writing – review and editing (equal); Funding acquisition (supporting)

**Marc T. J. Johnson:** Conceptualization (equal); Methodology (equal); Data curation (supporting); Formal analysis (supporting); Writing – review and editing (equal); Funding acquisition (lead)

## Conflict of Interest

The authors have no conflict of interest to disclose.

## Data Availability Statement

All data and R code are deposited on Zenodo (https://zenodo.org/doi/10.5281/zenodo.11038229; Murray-Stoker et al. 2024). Links to raster files will be included in the code and associated metadata, and the references for the rasters are provided in the main article and supplement. We highly encourage others to use the R code and to modify it as needed for their own projects, and we just ask that this article is cited in return for those who make use of our workflow.

## Notes

https://zenodo.org/records/11038230

### Competing Interest Statement

The authors have declared no competing interest.

