## Appendix S1 for "Comparing approaches to quantify urbanization on a multicontinental scale"

##### Supplemental Methods

We developed code to calculate 19 environmental variables spanning landcover, climate, and socioeconomic categories for all sites for each of the sampling designs (Table S1). Distance from the city centre was calculated for each site as the distance on an ellipsoid using the `geosphere` package (Hijmans 2019); we used the same coordinates for the city centres as the GLUE project (Santangelo et al. 2022). We estimated the impervious surface cover (ISC) using the 30-m resolution Global Man-Made Impervious Surface raster dataset (Brown de Colstoun et al. 2017), with ISC averaged within a 250-m buffer surrounding each site. We calculated the Human Influence Index (HII) for each site using the 1-km resolution HII raster dataset (Wildlife Conservation Society - WCS and Center for International Earth Science Information Network - CIESIN - Columbia University 2005). We downloaded the Normalized Difference Vegetation Index (NDVI) for each site across a 5-year period (1 January 2014-31 December 2018), which resulted in 115 measurements for each site (23 measurements per year). For each of the 5 years, we calculated the annual mean, minimum, and maximum NDVI and then averaged these estimates to generate a single mean, minimum, and maximum NDVI for each site; all NDVI measurements were calculated using the `MODISTools` package (Tuck et al. 2014). We calculated a suite of climate variables (mean annual temperature, temperature seasonality, range of annual temperature, annual precipitation, and precipitation seasonality) using the WorldClim database (Fick and Hijmans 2017) and 30 s resolution rasters. We also calculated an aridity index for each using ~1 km raster datasets from the CGIAR Consortium for Spatial Information (CGIAR-CSI 2019, Zomer et al. 2022).

We used globally-gridded estimates of gross domestic product (GDP) for 2005 and Shared Socioeconomic Pathways (SSPs) projections for years 2030 and 2100 using 1-km resolution raster files (Wang and Sun 2022). Each SSP is a projection based on a series of demographic, economic, and political development variables under global climate change, and each SSP has different constraints and assumptions for the projections. Briefly, SSP1 (sustainability) assumes a shift to more sustainable practices, increased environmental awareness, and a gradual shift to more sustainable and less resource-intensive lifestyles (O'Neill et al. 2014). Urbanization is expected to increase, as will human development (e.g., education, health investments, gender equality, equity). In contrast, SSP5 (fossil-fueled development) assumes fossil-fueled development and the maintenance of current policies on social and economic development (Dellink et al. 2017, O'Neill et al. 2017); urbanization is again expected to increase alongside human development. SSP2 (middle of the road) assumes current social and economic trends continue and do not strongly deviate from historical patterns, although with lower increases in urbanization coupled with lower improvements to human development ((Dellink et al. 2017, O'Neill et al. 2017). Values for environmental variables calculated from raster files (i.e., ISC, HII, climate variables, and socioeconomic variables) relied on custom functions using the `raster` package (Hijmans et al. 2023).

**Appendix S1**

#### References


Brown de Colstoun, E. C., C. Huang, P. Wang, J. C. Tilton, B. Tan, J. Phillips, S. Niemczura, P.-Y. Ling, and R. E. Wolfe. 2017. Global Man-made Impervious Surface (GMIS) Dataset From Landsat. NASA Socioeconomic Data and Applications Center (SEDAC), Palisades, New York.

CGIAR-CSI. 2019. Global Aridity Index and Potential Evapotranspiration Climate Database v3.

Dellink, R., J. Chateau, E. Lanzi, and B. Magné. 2017. Long-term economic growth projections in the Shared Socioeconomic Pathways. Global Environmental Change 42:200–214.

Fick, S. E., and R. J. Hijmans. 2017. WorldClim 2: new 1-km spatial resolution climate surfaces for global land areas. International Journal of Climatology 37:4302–4315.

Hijmans, R. J. 2019. geosphere: Spherical Trigonometry. <https://CRAN.R-project.org/package=geosphere>

Hijmans, R. J., J. van Etten, M. Sumner, J. Cheng, D. Baston, A. Bevan, R. Bivand, L. Busetto, M. Canty, B. Fasoli, D. Forrest, A. Ghosh, D. Golicher, J. Gray, J. A. Greenberg, P. Hiemstra, K. Hingee, A. Ilich, I. for M. A. Geosciences, C. Karney, M. Mattiuzzi, S. Mosher, B. Naimi, J. Nowosad, E. Pebesma, O. P. Lamigueiro, E. B. Racine, B. Rowlingson, A. Shortridge, B. Venables, and R. Wueest. 2023. raster: Geographic Data Analysis and Modeling. <https://cran.r-project.org/web/packages/raster/index.html>

O'Neill, B. C., E. Kriegler, K. L. Ebi, E. Kemp-Benedict, K. Riahi, D. S. Rothman, B. J. van Ruijven, D. P. van Vuuren, J. Birkmann, K. Kok, M. Levy, and W. Solecki. 2017. The roads ahead: Narratives for shared socioeconomic pathways describing world futures in the 21st century. Global Environmental Change 42:169–180.

O'Neill, B. C., E. Kriegler, K. Riahi, K. L. Ebi, S. Hallegatte, T. R. Carter, R. Mathur, and D. P. van Vuuren. 2014. A new scenario framework for climate change research: the concept of shared socioeconomic pathways. Climatic Change 122:387–400.

Santangelo, J. S., et al. 2022. Global urban environmental change drives adaptation in white clover. Science 375:1275–1281.

Tuck, S. L., H. R. P. Phillips, R. E. Hintzen, J. P. W. Scharlemann, A. Purvis, and L. N. Hudson. 2014. MODISTools - downloading and processing MODIS remotely sensed data in R. Ecology and Evolution 4:4658–4668.

Wang, T., and F. Sun. 2022. Global gridded GDP data set consistent with the shared socioeconomic pathways. Scientific Data 9:221.

Wildlife Conservation Society - WCS and Center for International Earth Science Information Network - CIESIN - Columbia University. 2005. Last of the Wild Project, Version 2, 2005 (LWP-2): Global Human Influence Index (HII) Dataset (Geographic). NASA Socioeconomic Data and Applications Center (SEDAC), Palisades, New York.

Zomer, R. J., J. Xu, and A. Trabucco. 2022. Version 3 of the Global Aridity Index and Potential Evapotranspiration Database. Scientific Data 9:409.

### Appendix S1

#### Supplemental Table

**Table S1:** List of environmental variables that were calculated for each site, along with the spatial resolution of the data source and brief description for each metric. Urbanization metrics were classified as landcover, climate, and socioeconomic. Variables are abbreviated as follows: ISC = impervious surface cover, HII = human influence index, NDVI = normalized difference vegetation index, GDP = gross domestic product, and SSP = shared socioeconomic pathway.

| Environmental Variable | Spatial Resolution | Classification | Description |
|---|---|---|---|
| **Distance** | Exact | Landcover | Distance from the city centre (km) |
| **ISC** | 30 m | Landcover | Mean impervious surface cover in a 250-m buffer surrounding each site (%) |
| **HII** | 1 km | Landcover | Composite measure of human influence, including population density, land use and infrastructure, and human access |
| **Mean NDVI** | 250 m | Landcover | Mean 'greenness' of the site; calculated within a 250-m buffer over a 5-year period |
| Min NDVI | 250 m | Landcover | Minimum 'greenness' of the site; calculated within a 250-m buffer over a 5-year period |
| Max NDVI | 250 m | Landcover | Maximum 'greenness' of the site; calculated within a 250-m buffer over a 5-year period |
| **Mean Annual Temperature** | 30 s (~1 km) | Climate | Mean local air temperature (°C) |
| Temperature Seasonality | 30 s (~1 km) | Climate | Variation in local temperature (standard deviation, °C) |
| Range Annual Temperature | 30 s (~1 km) | Climate | Range in local air temperature (°C) |
| **Annual Precipitation** | 30 s (~1 km) | Climate | Local precipitation (mm) |
| Precipitation Seasonality | 30 s (~1 km) | Climate | Variation in local precipitation (coefficient of variation) |
| Aridity Index | 1 km | Climate | Balance between precipitation and evapotranspiration |
| GDP 2005 | 30 s (~1 km) | Socioeconomic | Gross domestic product for 2005 (United States dollars) |
| SSP1 2030 | 30 s (~1 km) | Socioeconomic | GDP under SSP1 (sustainability) extrapolated to 2030 (USD) |
| SSP1 2100 | 30 s (~1 km) | Socioeconomic | GDP under SSP1 (sustainability) extrapolated to 2100 (USD) |
| **SSP2 2030** | 30 s (~1 km) | Socioeconomic | GDP under SSP2 (middle of the road) extrapolated to 2030 (USD) |
| **SSP2 2100** | 30 s (~1 km) | Socioeconomic | GDP under SSP2 (middle of the road) extrapolated to 2100 (USD) |
| SSP5 2030 | 30 s (~1 km) | Socioeconomic | GDP under SSP5 (fossil-fueled development) extrapolated to 2030 (USD) |
| SSP5 2100 | 30 s (~1 km) | Socioeconomic | GDP under SSP5 (fossil-fueled development) extrapolated to 2100 (USD) |

Note: Focal environmental variables are in bold.

### Appendix S1

#### Supplemental Figures

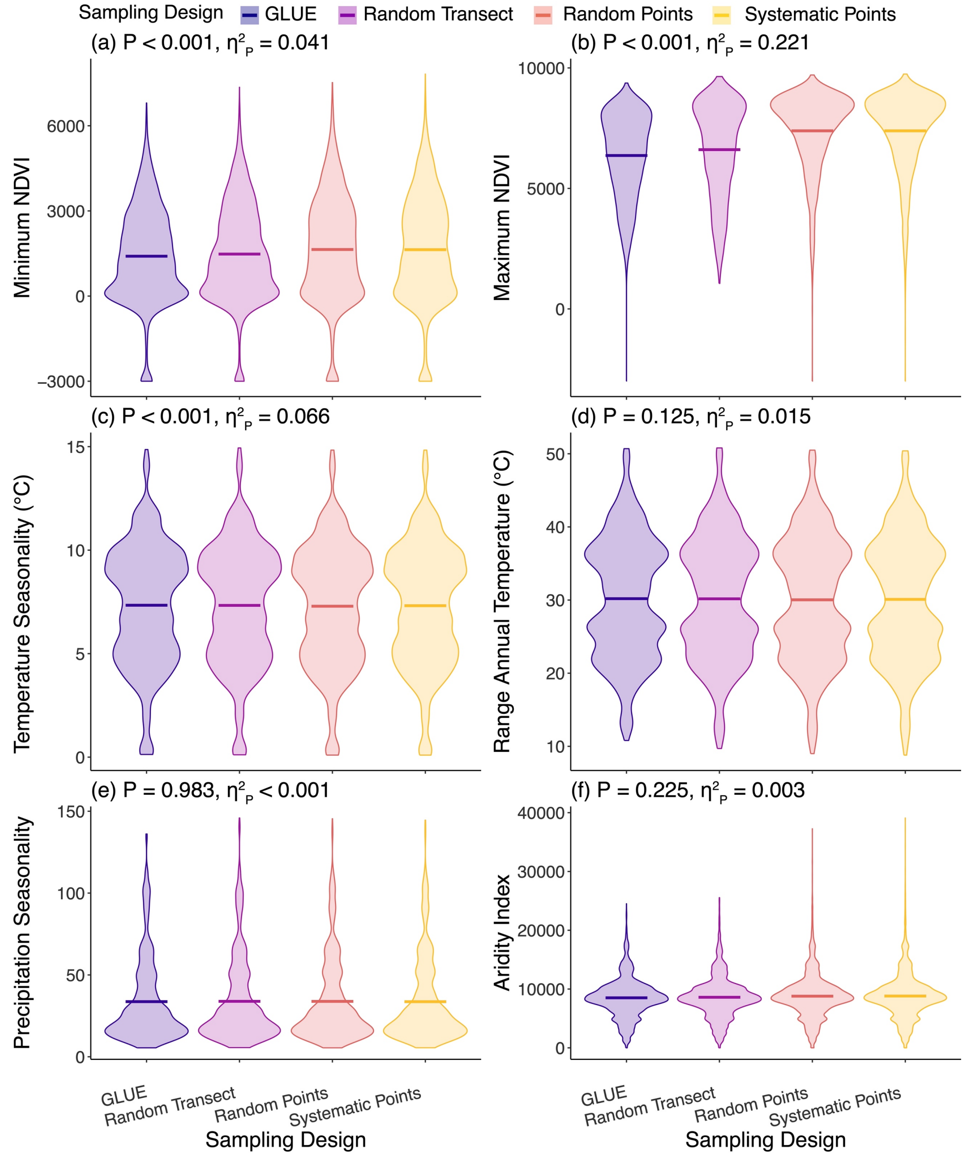


**Figure S1:** Violin plots of non-focal landcover and climate variables by sampling design. Violins are colored by sampling design, where GLUE transect = dark blue, random transect = purple, random points = orange, and systematic points = yellow. Crossbars indicate the mean. Inset text provides the P-value and effect size for the fixed effect of sampling design (partial eta-squared, $\eta^2_P$). Detailed test statistics are provided in Table 1.

#### Appendix S1

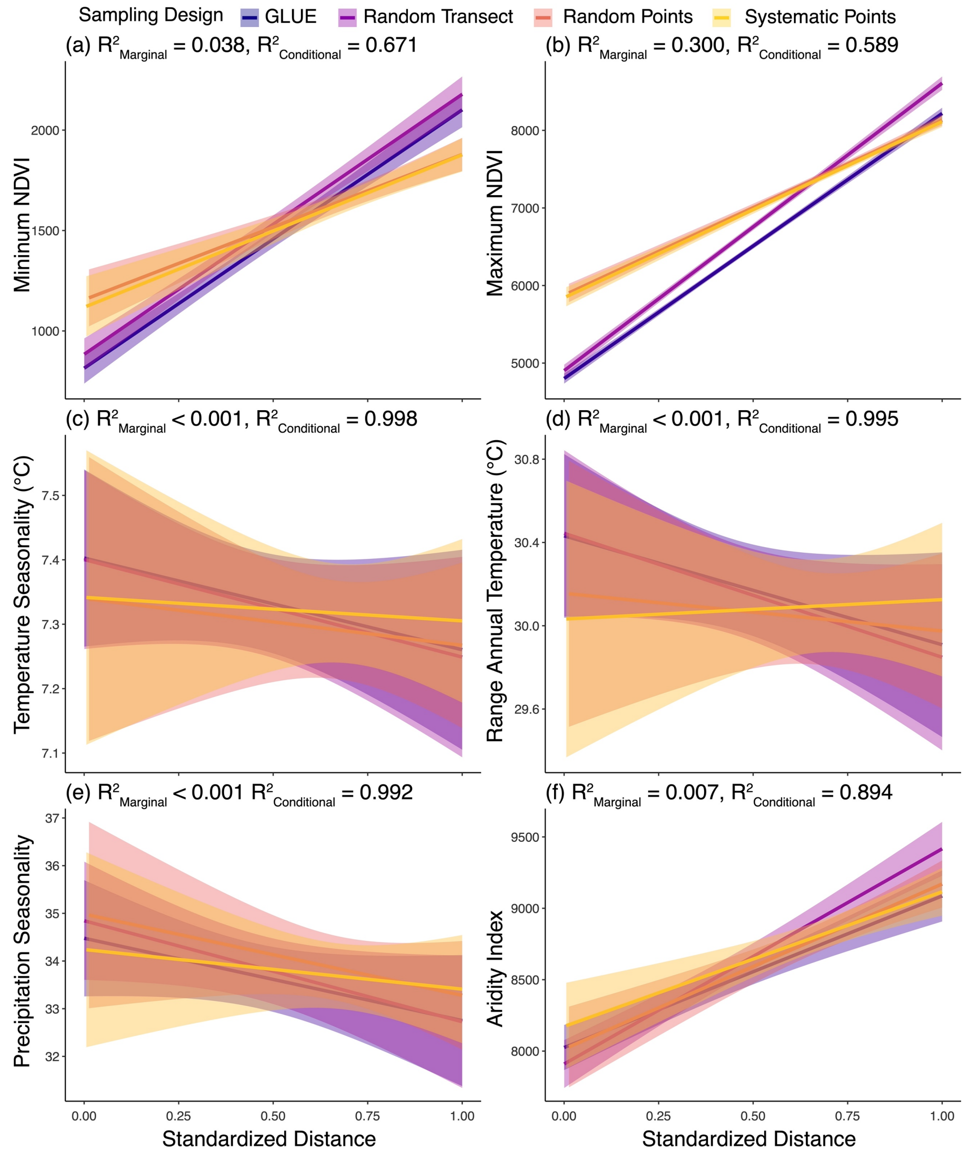


**Figure S2:** Relationships between non-focal environmental variables (landcover and climate) and standardized distance to the city centre (0 = city centre, 1 = rural limit) by sampling design. Lines are lines-of-best fit (± 95% confidence interval) and are coloured by sampling design: GLUE transect = dark blue, random transect = purple, random points = orange, and systematic points = yellow. We report the marginal $R^2$ (variance explained by fixed effects, $R^2_{\text{Marginal}}$) and conditional $R^2$ (variance explained by fixed and random effects, $R^2_{\text{Conditional}}$). Detailed test statistics provided in Table 1.

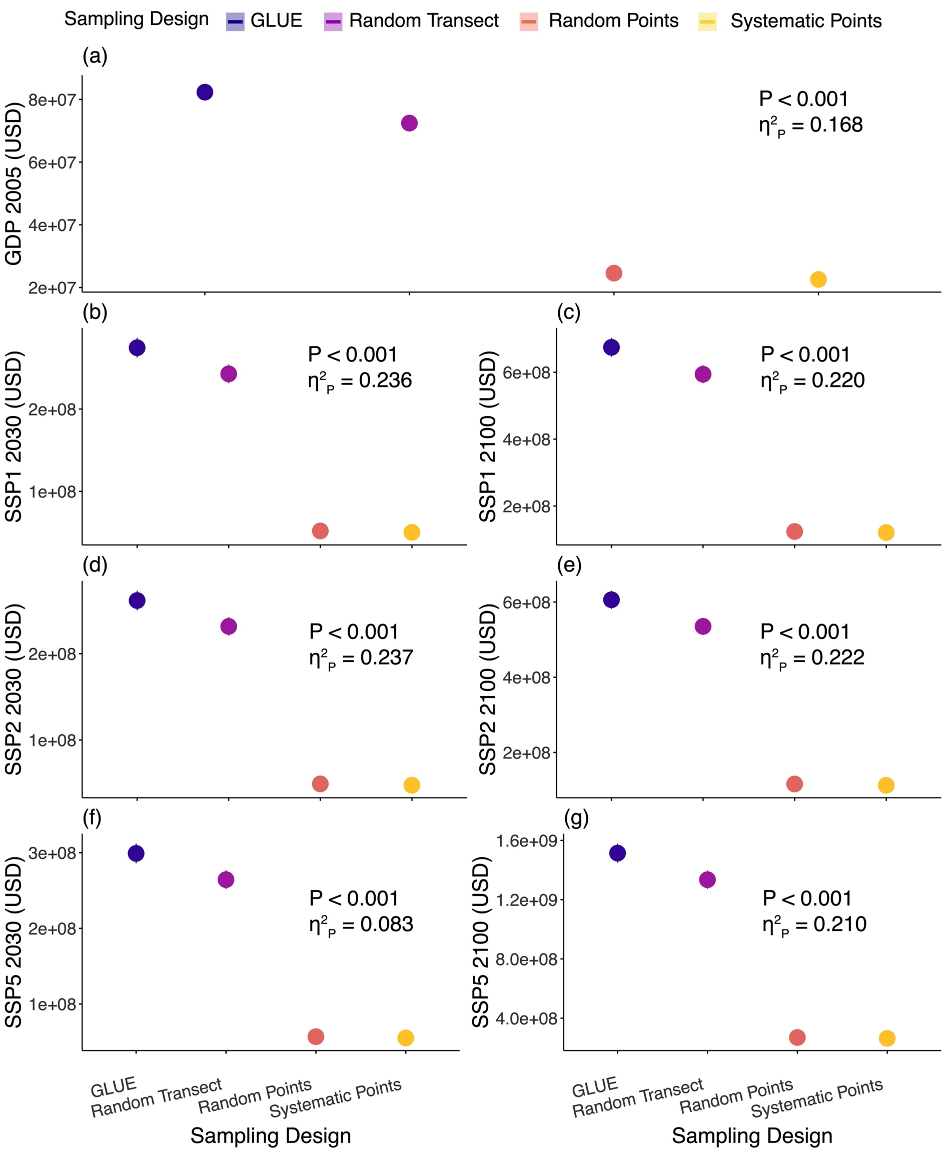


**Figure S3:** Estimates for socioeconomic variables (mean ± SE) by sampling design. Points are coloured by sampling design, where GLUE transect = dark blue, random transect = purple, random points = orange, and systematic points = yellow. Inset text provides the P-value and effect size for the fixed effect of sampling design (partial eta-squared, $\eta^2_P$). Detailed test statistics are provided in Table 1. Variables are abbreviated as GDP = gross domestic product and SSP = shared socioeconomic pathway.

### Appendix S1

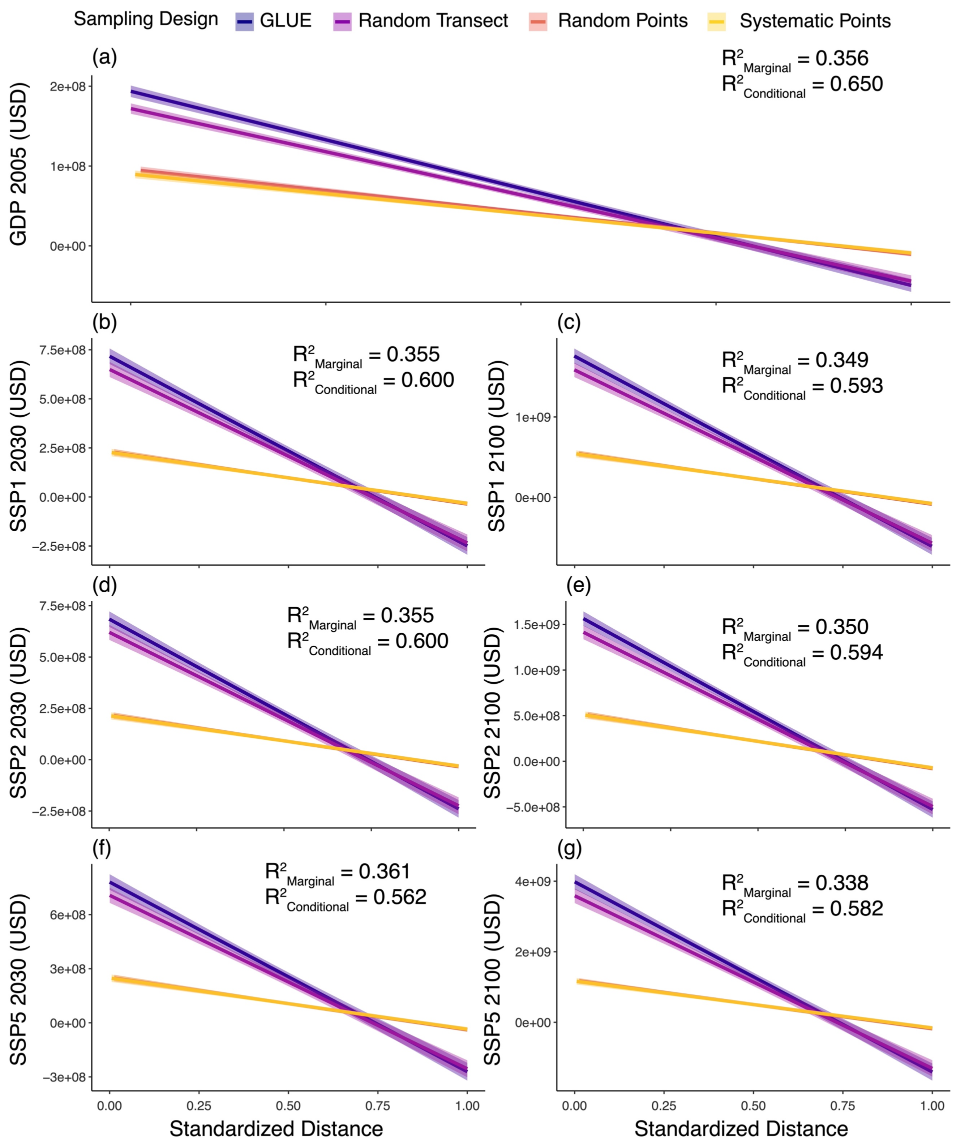


**Figure S4:** Relationships between socioeconomic variables and standardized distance to the city centre (0 = city centre, 1 = rural limit) by sampling design. Lines are lines-of-best fit (± 95% confidence interval) and are coloured by sampling design: GLUE transect = dark blue, random transect = purple, random points = orange, and systematic points = yellow. We report the marginal $R^2$ (variance explained by fixed effects, $R^2_{Marginal}$) and conditional $R^2$ (variance explained by fixed and random effects, $R^2_{Conditional}$). Detailed test statistics provided in Table 1. Variables are abbreviated as GDP = gross domestic product and SSP = shared socioeconomic pathway.

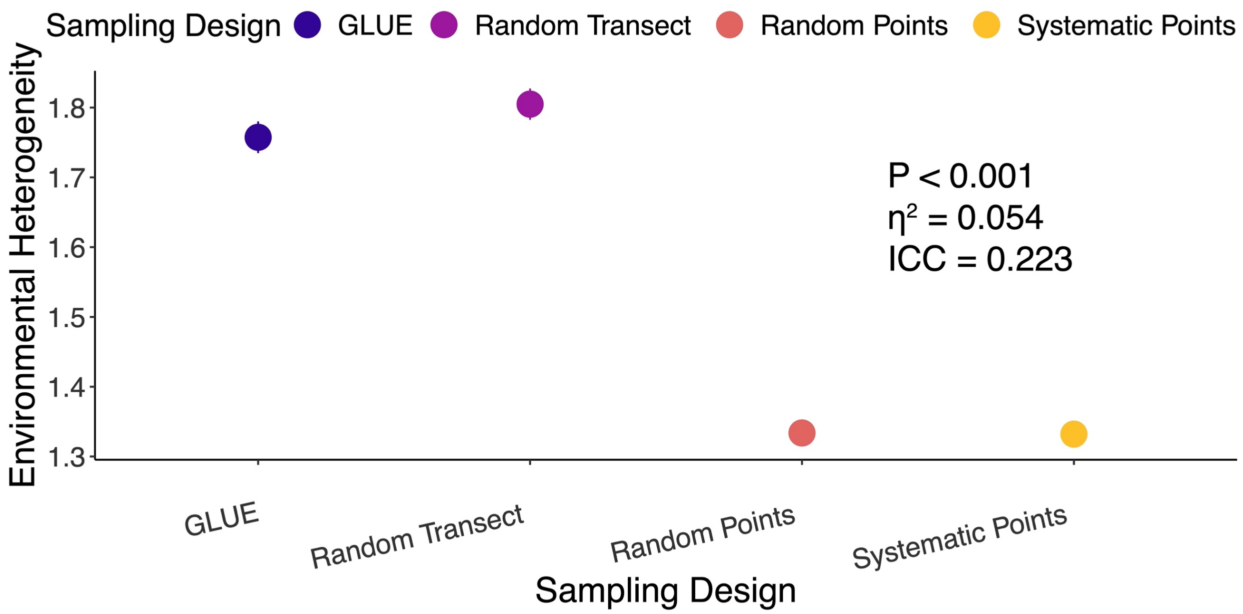


**Figure S5:** Environmental heterogeneity (mean ± SE) by sampling design. Points are colored by sampling design, where GLUE transect = dark blue, random transect = purple, random points = orange, and systematic points = yellow. Inset text provides the P-value and effect size for the fixed effect of city (eta-squared, $\eta^2$) from the ANOVA. We also report the effect size for the random effect of city (intraclass correlation coefficient, ICC).

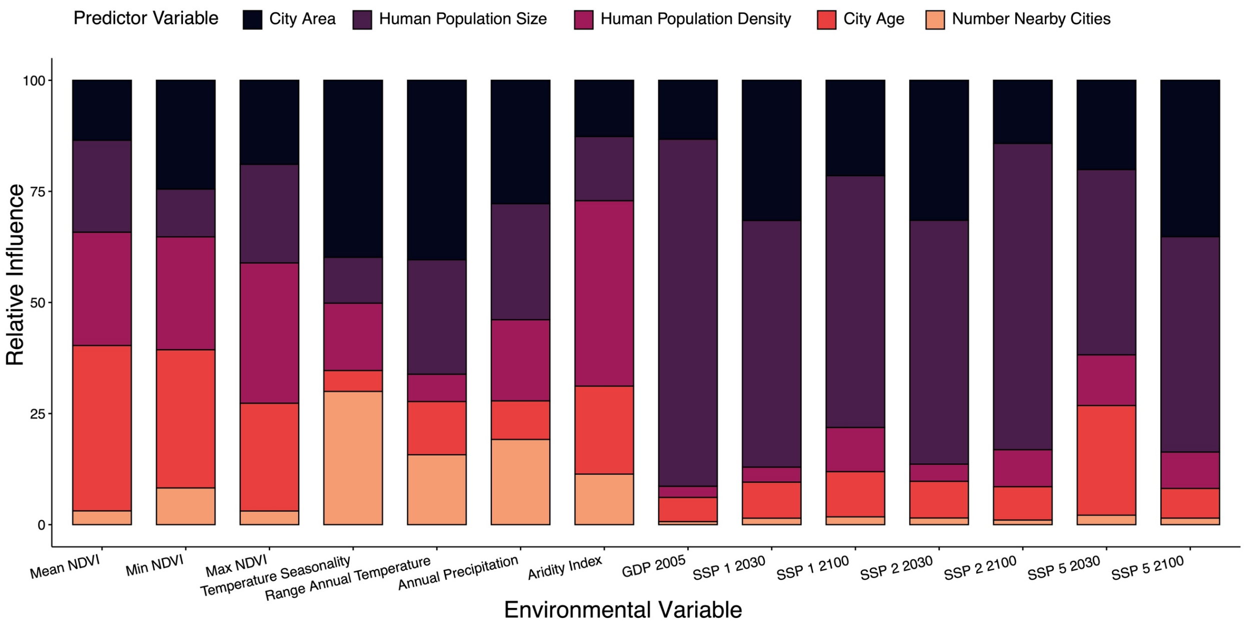


**Figure S6:** Relative influence of city characteristics (city area, human population size, human population density, city age, and number of nearby cities) for predicting deviations between transects (GLUE and random transects) and points (random and systematic points) for non-focal landcover, climate, and socioeconomic variables. Variables are abbreviated as: NDVI = normalized difference vegetation index, GDP = gross domestic product, and SSP = shared socioeconomic pathway.
